# An Open-Source Plate Reader

**DOI:** 10.1101/413781

**Authors:** Karol Szymula, Michael S. Magaraci, Michael Patterson, Andrew Clark, Sevile G. Mannickarottu, Brian Y. Chow

## Abstract

Microplate readers are foundational instruments in experimental biology and bioengineering that enable multiplexed spectrophotometric measurements. To enhance their accessibility, we here report the design, construction, validation, and benchmarking of an open-source microplate reader. The system features full-spectrum absorbance and fluorescence emission detection, *in situ* optogenetic stimulation, and stand-alone touch screen programming of automated assay protocols. The total system costs <$3500, a fraction of the cost of commercial plate readers, and can detect the fluorescence of common dyes down to ∼10 nanomolar concentration. Functional capabilities were demonstrated in context of synthetic biology, optogenetics, and photosensory biology: by steady-state measurements of ligand-induced reporter gene expression in a model of bacterial quorum sensing, and by flavin photocycling kinetic measurements of a LOV (light-oxygen-voltage) domain photoreceptor used for optogenetic transcriptional activation. Fully detailed guides for assembling the device and automating it using the custom Python-based API (Application Program Interface) are provided. This work contributes a key technology to the growing community-wide infrastructure of open-source biology-focused hardware, whose creation is facilitated by rapid prototyping capabilities and low-cost electronics, optoelectronics, and microcomputers.

**Table of Contents Graphic:** 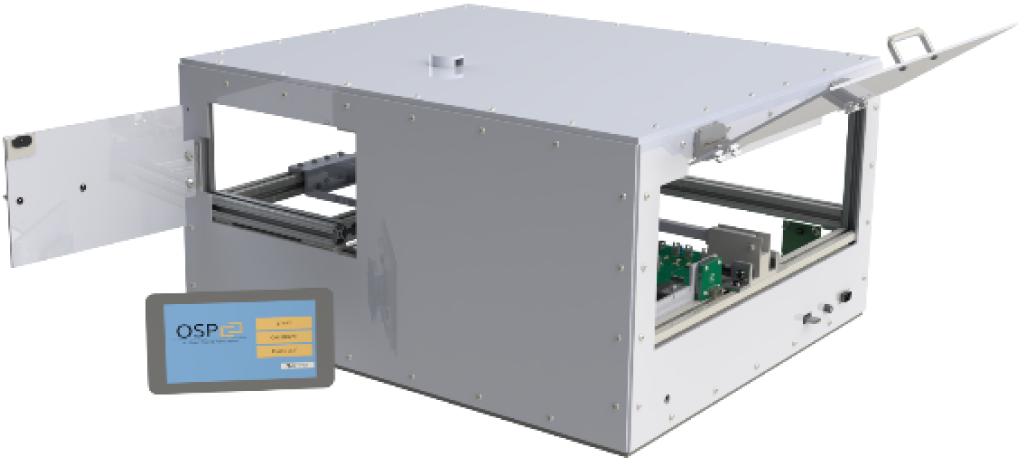

## Introduction

The creation of open-source hardware for enabling molecular and cellular studies is facilitated by the ready availability of rapid prototyping techniques, low-cost opto-electronics, and commoditized microcomputers & microcontrollers. Such open-source platforms - which to date include fluorescence imagers ^1-2^, spectrophotometers ^3^, turbidostats ^4^, robotic liquid handlers ^5-6^, and multi-well plate illuminators for optogenetic stimulation ^7-8^ - increase access to both standard and custom laboratory apparatus alike. Here, we report the creation of an open-source plate reader (OSP), which is one of the most ubiquitous and important technologies in experimental biochemistry, biology, and bioengineering, because a plate reader permits automated and multiplexed spectrophotometric measurements.

Plate readers typically consist of a multi-mode (absorbance / optical density, and fluorescence) spectrophotometer, plus a programmable-motion stage for multiplexing by serial positioning of sample wells of the plate over a single detector. Detection wavelengths are selected using diffraction gratings or dichroic filters, and excitation wavelengths are selected similarly or by using monochromatic or narrow-band excitation sources such as lasers and light-emitting diodes (LEDs). They may possess auxiliary input ports for fluid delivery, and these ports have been successfully co-opted for other purposes such as optogenetic stimulation ^9^. However, commercial plate readers cost tens of thousands of $USD, which can be prohibitive for resource-limited environments including educational settings.

The open-source framework (**Figure 1**) reported here is intended to provide the necessary and modular functionalities of a commercial system, while maintaining the performance required for most assays at a fractional cost (<$3500 or ∼10x reduction). The system features full visible spectrum absorbance and fluorescence emission capabilities, *in situ* optogenetic stimulation, and stand-alone touch screen programming of automated assay protocols. It is expandable and customizable, with built-in access- and communication ports for external modules to enhance function and its own Python-based Application Programming Interface (API). We document the design considerations, overview of system assembly and operation, benchmarking with standard fluorochrome dyes, and validation in molecular and cellular-level experiments in synthetic biology, optogenetics, and photobiology contexts.

**Figure 1.**
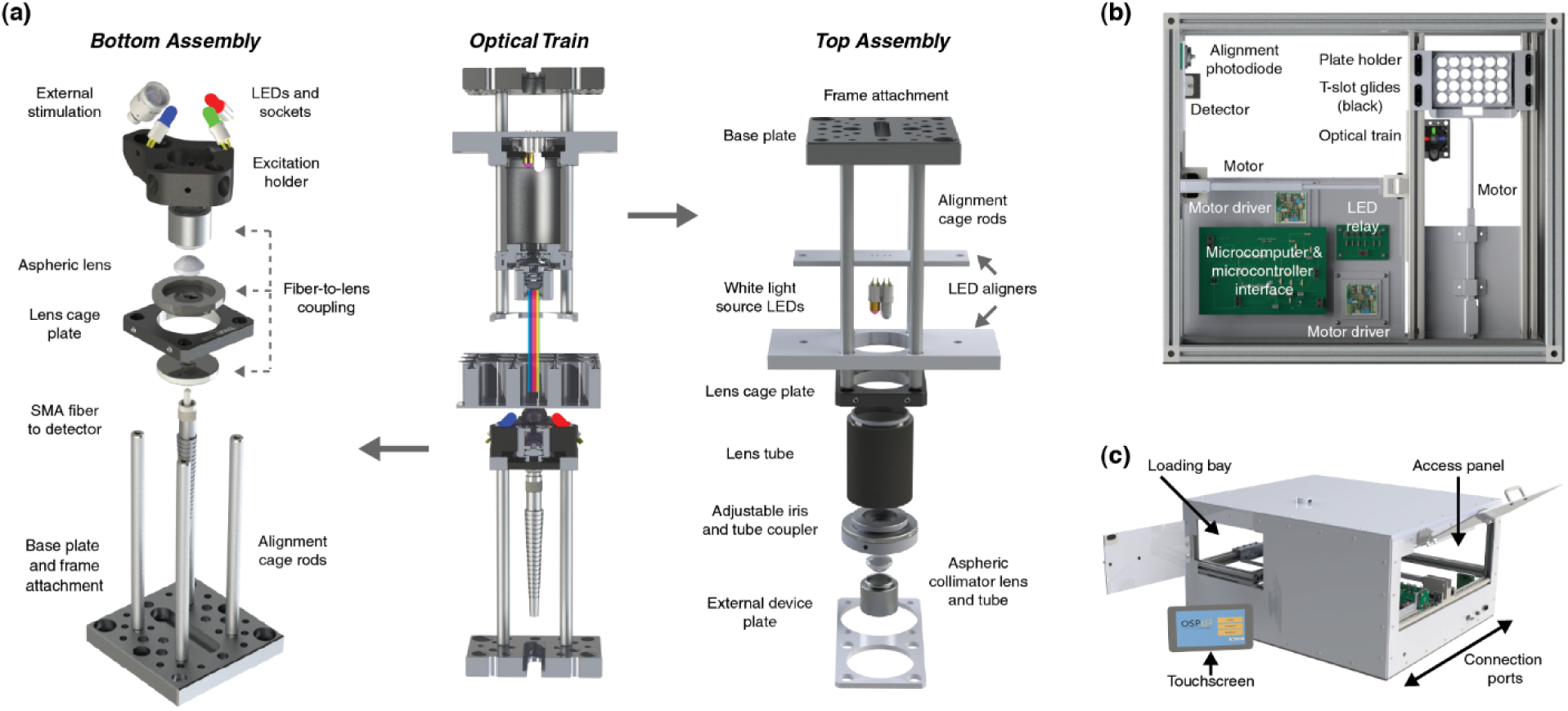
Opto-mechanical summary of the open-source plate reader (OSP). All images are CAD renderings, **(a)** The optical train, with exploded views of bottom assembly for detection and optical excitation, and the top assembly for the white light source for optical density measurements and for external devices (e.g. high-power optogenetic stimulation or fluid delivery). The white light source combines two LEDs to span the visible spectrum, **(b)** Top-down view of the whole instrument. The relay board contains header blocks that consolidate connections of the individual device components, separated from the main microcomputer board. The top optical train is omitted for clarity, **(c)** Isometric view of the instrument, with touchscreen that can be mounted. All connections and access for repair are consolidated on one side.

## Results and Discussion

### Opto-mechanical platform design

The fundamental core of a plate reader is two-axis motion for x-y positioning with respect to a single set of fiber optic input/outputs, which enables multiplexing. Thus, we designed low-cost hardware and software frameworks (**Figures 1 and 2**) for automated motion, and for interfacing with open-source devices and fiber-coupled optoelectronics. For baseline functionality useful in most laboratory contexts, the open-source plate reader (OSP) employs a CCD (charge-coupled device) camera-based detector with open-source drivers (Ocean Optics), and inexpensive dome-capped light-emitting diodes (LEDs) as illumination sources for absorbance / optical density (Od) measurements and for fluorescence measurements with common fluorochromes (**Figure 1a**). The use of fiber optics to couple the optical elements to the sampling location allows for modular choice in excitation sources and detectors, and importantly, also supports the expansion or customization through these optical fibers to more sensitive detectors or to sources of higher power or different wavelengths. Thus, the system is designed as both a low-cost and expandable plate reader chassis.

**Figure 2.**
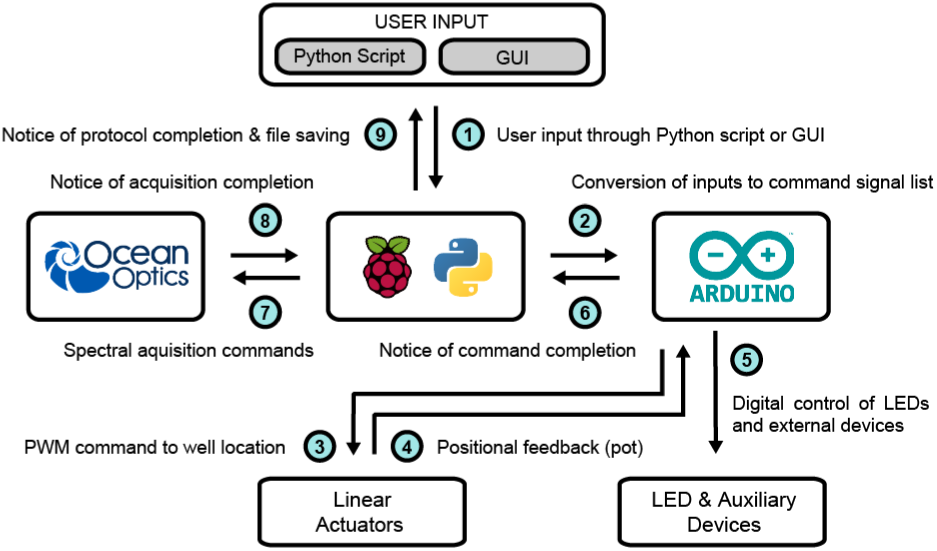
Device integration and automation workflow. User inputs to a Raspberry Pi microcomputer (center) are entered via the graphical user interface (GUI) or written Python script. The microcomputer coordinates the inputs and outputs to all other components. PWM = pulse width modulation, pot = potentiometer value. LED = light-emitting diode.

For automated motion, a custom 3D-printed plate holder is attached to low-cost linear drive motors that provide positional feedback voltage readings. The motors were chosen to ensure sufficient travel range (140 mm) for plate loading and load-bearing capacity (100 N) for the plate stage. The motor-connected stage glides on a modular frame of extruded t-slotted aluminum, which was chosen for its ease of assembly and utility of the t-slot tracks themselves for alignment, similar to dovetail rails (**Figure 1b**). Plates are tightly and repeatably positioned into one corner of the holder by spring-loading, and loaded through a front door (**Figure 1c**).

For optical detection, an optomechanical lens cage assembly under the plate holds a collimating aspheric lens to couple transmitted and emitted (bottom fluorescence) light to a multimode fiber optic cable (**Figure 1a, *left*).** This patch cord can be connected to any fiber-compatible detector with a SMA fitting. In the baseline low-cost version here, we use a camera-based Ocean Optics visible-range detector because of its available open-source drivers and overall balance of sensitivity, full-spectrum- and monochromatic detection, and relative low-cost compared to monochromator-coupled photomultiplier tubes (PMT).

For bottom fluorescence excitation, a 3D-printed component surrounds the detector fiber input assembly, and projects multiple light-sources onto the plate bottom. Three positions hold sockets connectors for dome-capped LEDs as cost-effective, bright, and built-in fluorescence excitation sources. Initially, these LEDs are blue, green, and orange (λ = 470 nm, 525 nm, and 610 nm) for exciting common dyes and fluorescent proteins, but they can be simply replaced with different standard dome-capped LEDs. *In lieu* of using collimating lenses, the LEDs have narrow solid angles which, combined with the acceptance angle of the collection aspheric, prevents well-to-well crosstalk. These LEDs can also be used for optogenetic stimulation. The OSP was designed for visible spectrum measurements of most plate reader assays, and thus similar to any commercial filter-based plate reader, omits a fully tunable excitation source like a scanning monochromator for fluorescence excitation scans.

An additional position of the projection assembly holds a SMA-terminated collimated fiber optic for use with external sources as alternatives or upgrades to dome-capped LEDs. For cases where the user requires multiple external sources coupled to the one collimated projection fiber, we suggest commercially available bifurcated or fan-out optical fibers (e.g. from Thorlabs) as a cost-effective alternative to traditional dichroic beam combiners, because the fiber-based solution is effectively universal by eliminating the minimum spectral separation for co-aligning optical sources with dichroic mirrors. However, such combiners are not recommended for co-aligning a detection fiber with excitation sources (i.e. 90° incident angle between sample and excitation), because direct back reflection from a collimating lens or plate bottom saturates the detector in the absence of an automated dichroic filter wheel or tunable diffraction gratings.

For top-side excitation for absorbance / optical density measurements and optogenetic stimulation, another optomechanical lens train (**Figure 1a, *right***) over the plate positions a visible-spectrum white light source, consisting of two LEDs (LED 1: λ = 430 - 660 nm, LED2: λ = 430 nm) (**Supplementary Figure 1**). A laser-cut component connected to the top optical train positions an auxiliary port for external functionality like fluid delivery and optogenetic stimulation. Similar to commercial systems, the auxiliary port is offset from the detector with a ∼5 sec delay between delivery of fluid/photons to the well and measurement from the well, although simultaneous of zero-delay optogenetic stimulation is possible using the bottom excitation sources. The auxiliary port is designed to couple to a standard optical lens cage plate for attaching and aligning custom components, which can be fed into the OSP through a grommet on top.

### Automation and Interface Design

The OSP can be operated by an external computer or Raspberry Pi microcomputer by using automated scripts or a Graphical User Interface (GUI) that supports touchscreen control (**Figure 2**). The GUI and automation software are programmed in Python. The description of the interface will assume the use of a Raspberry Pi microcomputer from hereon for simplicity.

The Raspberry Pi communicates directly with the detector to set acquisition parameters, and also sends programming commands to an Arduino microcontroller (**Supplementary Figure 2 and Supplementary Note 1**). The Arduino controls the other functions (LEDs, motors, and auxiliary functions) and synchronizes them with the detector data acquisition through digital logic. LED timing is controlled by digital switching of a transistor-gated current source. The motors are controlled by pulse width modulation commands sent to the commercial motor driver-boards, which also relay positional feedback (potentiometer readings). Auxiliary functions can be triggered by standard digital logic control of external devices.

The GUI (**Supplementary Note 1**) emulates the basic programming capabilities of commercial plate readers. Acquisition parameters include: (i) detector integration time, (ii) selected data range for spectra (scan averaging, boxcar, excitation wavelength), (iii) time-step for kinetic acquisition, (iv) well selection and control (ordering, calibration (**Supplementary Figure 3**)), and (v) coordination with auxiliary functions. Data spreadsheets are exported to local storage and can be manipulated locally in the Raspberry Pi environment using the open-source office software suite, LibreOffice (or similar).

**Figure 3.**
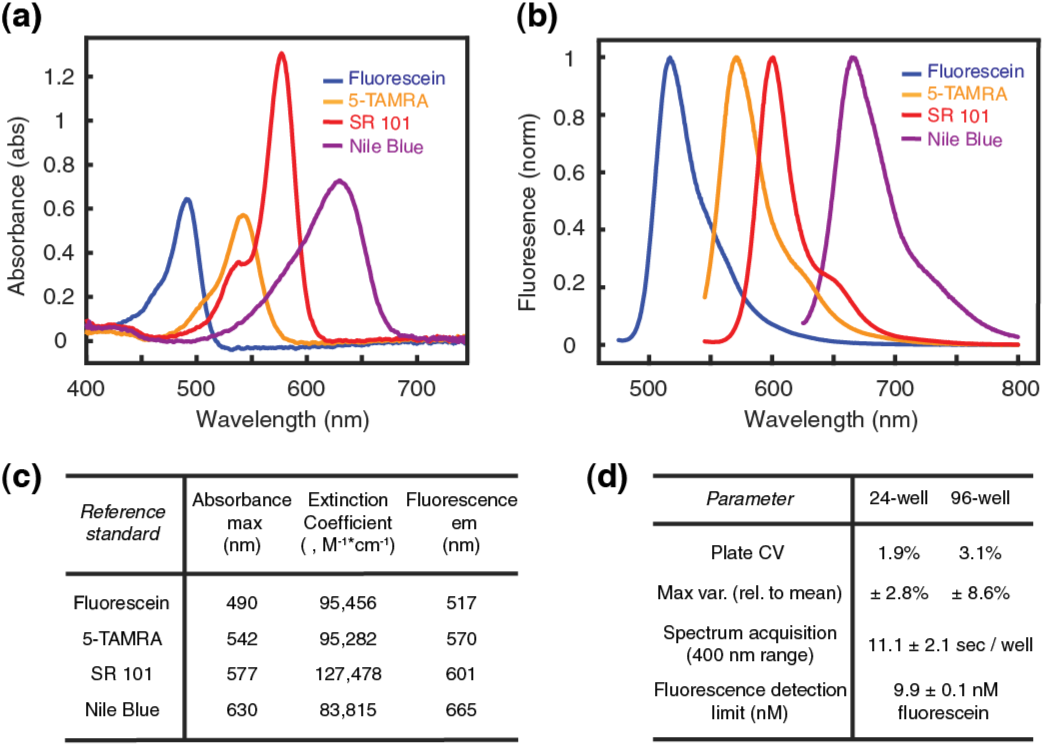
Characterization of the OSP using reference dyes. Measurements were made according to commercial protocols (Thermo R14872). 5-TAMRA = 5-carboxytetramethylrhodamine; SR 101 = Sulforhodamine 101. **(a)** Absorbance spectra, **(b)** Fluorescence emission spectrum, **(c)** Key measured spectral parameters from a-b. **(d)** Key performance ranges. CV = Coefficient of variation across plate. Max var. = Variation range across plate. Spectrum acquisition time with 1.25 sec integration and includes stage motion to well. Detection limit error = stdev.

### Protocols: Device Construction and Operation

Detailed step-by-step assembly guides and programming overview are available (**Supplementary Notes 1-4**), as summarized here. Supplementary Note 1 describes software installation, the organization of the open-source API, the available functions for custom programming, and a walk-through of the GUI operation. Supplementary Note 2 provides assembly instructions for the top and bottom optical trains shown in Figure 1a. Supplementary Note 3 describes the assembly of the printed circuit boards and the photodiode assembly used for calibrating the spatial coordinates of the motor. Supplementary Note 4 provides overall assembly instructions of the t-slotted frame, all custom 3D-printed or laser cut components including the plate stage (**Supplementary File 1**), and the integration of all subcomponents (trains, circuit boards, stage, and frame) into a complete device. The total cost of the system is <$3,500 (**Supplementary Table 1**).

The use of lens cage assemblies allows for facile alignment of all optical components along the detection axis (**Supplementary Note 3**). To initialize the spatial coordinates of the microplate wells post-assembly (**Supplementary Figure 3**), transmission measurements are made through a mask overlaid on the plate, as detected by a fast photodiode (**Figure 1b**); an automated script identifies the center of three alignment wells and the plate edges to define the spatial map. This calibration protocol takes ∼3 minutes, which is short enough to regularly perform to ensure repeatability.

An .*iso* file for software installation (*will be*) available via a permanent link at the University of Pennsylvania’s Scholarly Commons, where all necessary guides, schematics, software, and CAD files will be updated with version improvements; this site (*will be*) mirrored at the GitHub software repository.

### Performance Benchmarking with Reference Dyes

The baseline system performance was assessed using a reference standard fluorescent dye kit (**Figure 3**). The respective absorbance/emission peak spectra and extinction coefficients of the dyes were in line with expected values. Thus, the custom opto-mechanical elements do not introduce major aberrations. The fluorescence of NIST-traceable fluorescein could be measured with a ∼10 nM limit of detection with the low-cost detector. Such a detection limit is sufficient for common fluorescence assays as intended, albeit greater than the ∼10 pM limit of commercial systems that use PMTs.

The spectral separation required between fluorescence excitation and emission detection starting-wavelength was approximately one FWHM (full-width at half-maximum) of an LED because of the collection of reflected light from the plate bottom (**Supplementary Figure 4**). Given a typical ∼30nm FWHM of dome-capped LEDs here, the separation is on par with the recommended separation of a commercial plate reader. The separation can be decreased as low-cost LEDs with narrower bandwidths become available, or with laser diodes.

**Figure 4.**
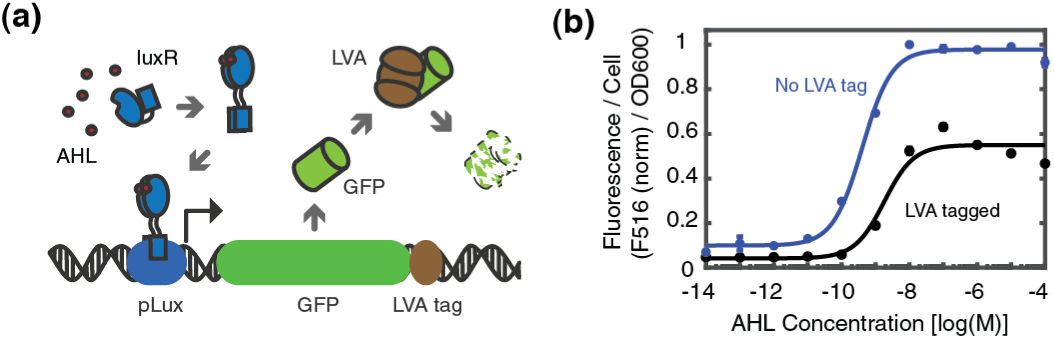
Cellular assays of a model of bacterial quorum sensing, **(a)** Schematized genetic circuit of GFP expressed under the transcriptional regulation of the **AHL** ligand-inducible luxR transcriptional activator, and under the post-translational degradatory regulation of **LVA** protease, **(b)** Transfer function between the **AHL** input and GFP fluorescence output, with and without the **LVA** tag. Switching point values agree with previous reports [Ref **1],** F516 = Fluorescence at λ = 516nm, scaled to max. OD6OO = Optical density at λ = 600nm. Error = s.e.m. (N=3).

**Figure 5.**
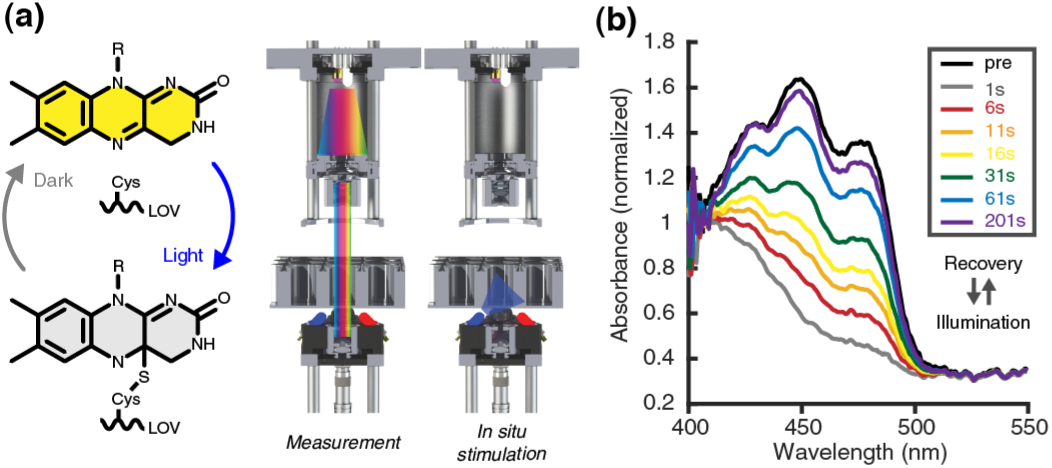
Flavin photocycling of EL222 by in-situ optogenetic stimulation. **(a)** Flavin photoadduct formation/reversion in a LOV sensor domain, measured by absorbance spectroscopy with in-situ stimulation (λ = 470 nm @ 10 mW/cm2, 5 sec), **(b)** Representative flavin photocycling of 37 µ M FPLC-purified recombinant holoprotein, normalized to its isobestic point. The triplet peak vibronic structure could be resolved, and the measured thermal reversion kinetics (τ = 31.5 sec, Cl = ± 0.5 sec, N = 3; 5 sec sampling interval, measured at λ *=* 450nm) were in agreement with reported values. Pre = Pre-optogenetic stimulation, taken in the dark. Times = Post-illumination recovery in the dark.

The coefficient of variations across entire 24-well or 96-well plates were 1.9% and 3.1%, respectively, with maximum variation from the mean of 2.8% and 8.6%, respectively. The variability is largely set by the motor resolution (vs. that of high-precision stepper motors that are ∼50-fold more expensive). The precision is sufficient for these common 24-well and 96-well plate formats, albeit unlikely for higher density 384-well plates and beyond.

The OSP moves acquires a full spectrum in ∼11 sec per well (inclusive of the movement to the well). Because the CCD detector captures all wavelengths simultaneously, the acquisition time for a full spectrum is the same as a single wavelength measurement. While a commercial plate reader with a PMT detects single wavelength in roughly the same time, the latter is much slower when measuring full spectra due to the need to sweep individual wavelengths; as a matter of reference, in our hands a Tecan microplate reader takes nearly six minutes to acquire a spectrum (400 nm-wide range with 2 nm step-size). This difference in time can be meaningful with samples in organic solvents prone to evaporation, or samples prone to sedimentation.

### Validation in Cellular and Biomolecular Contexts

We performed cellular and protein-level assays to demonstrate the standard and non-standard measurement functionalities of the OSP in biomolecular contexts. First, we measured the input/output transfer functions of bacterial “Receiver” strains ^1,10^, where a fluorescent protein reporter expression is placed under the switchlike transcriptional regulation of the (acyl-homoserine-lactone) AHL-responsive *luxR* activator (Figure 4). This system was chosen for its widespread use in synthetic biology to systematically study transcriptional regulation and inter-cellular communication ^10-11^, and also because it involves the two major functionalities of a plate reader of transmission measurements (optical density, absorbance) and fluorescence measurements needed to determine OD-normalized cellular fluorescence. The two *E. coli* strains differ by the presence or absence of an LVA degradation tag ^12-13^ on the GFP reporter. The relative switching behaviors of these strains are consistent with our previously reported measurements on a commercial (Tecan) plate reader (see BC-A1-001 and BC-A1-002 in Supplement of Reference: ^1^) with respect to half-saturation concentration of AHL and the relative magnitudes of GFP expression.

Next, we measured the flavin photocycling ^14^ kinetics of the light-oxygen-voltage domain (LOV) photoreceptor EL222 from purified recombinant protein ^15^. This kinetic measurement (absorbance at 450nm, A450) monitors the blue light-induced formation and thermal reversion (in the dark) of a cysteinyl-flavin photoadduct that underlies photosensory signaling by LOV proteins, which are of great importance in photobiology and pervasive in use in optogenetics and synthetic biology ^16-17^. EL222 in particular is commonly studied to explore the structure-function of light-induced homodimerization, and the bacterial LOV is useful for light-activated transcriptional activation with high on/off contrast ratios in eukaryotes ^18-19^. The photocycling measurement demonstrates the ability to measure a kinetic time-course and coordinate the measurement with a programmed set of events, in this case, *in situ* optogenetic stimulation of an individual well. To date, such *in situ* illumination for multiplexed measurements requires custom modification of a commercial plate reader ^9^.

In summary, we report the design, construction, and validation of an open-source plate reader framework. The cost-effectiveness of the OSP highlights the importance to laboratory research of rapid prototyping technologies, low-cost solid-state opto-electronics, and minimal single-board micro-computers and micro-controllers. The open-source nature and modularity of the system will facilitate future customization or enhancements that incorporate improved components as they decrease in cost, such as high-power collimated LEDs and the use of metal 3D-printing. The low-cost, modular, and expandable plate reader will enhance access to, and flexibility of, a core instrument across biological and chemical disciplines.

## Materials and Methods

The parts list, step-by-step guides for hardware construction and calibration, and software installation and application programming can be found in Supplementary Material, as described in “Protocols: Device Construction and Operation.”

All assays were cross-validated on a commercial Tecan Infinite M200 plate reader. The volume-dependent effective path length in 24-well plates was calculated from Beer’s Law plots of NIST-traceable fluorescein (F36915, NIST Standard Reference Material 1932 ^20^) at A(490 nm) assuming ε = 87,000 M^-1^ cm^-1^ [Conversion factor: 2.18 mL fluid ∼ 1 cm effective pathlength, linear down to 500 µL volume]. Reference dye measurements were performed according to kit manufacturer protocols (Thermo Fisher R14782), using an integration period of 1.25 seconds. Quorum sensing experiments were performed as described previously ^1^. Recombinant EL222 was bacterially produced and FPLC-purified as described previously ^21^. Fluorescence detection limit was measured by reported methods for determining plate reader sensitivity ^22^, with 0.3 µM NIST-traceable fluorescein and an integration time of 1.25 seconds.

## Supplementary Materials

[Tables] *Supplementary Table 1:* Parts list and cost summary. [Notes and Files] *Supplementary Note 1:* Software guide. *Supplementary Note 2:* Optical train subcomponent assembly guide. *Supplementary Note 3:* PCB and photodiode sub-component assembly guide. *Supplementary Note 4:* General assembly guide. *Supplementary File 1:* Zip file containing CAD files for all custom components and adapters. Future updates will be posted at the University of Pennsylvania Scholarly Commons, through a permanent link. [Figures] *Supplementary Figure 1:* White light-source spectrum. *Supplementary Figure 2:* API workflow. *Supplementary Figure 3:* Calibration of plate spatial coordinates. *Supplementary 4:* Fluorescence spectra without longpass removal of excitation.

## Author Contributions

KS and MSM constructed the device and application programming interface, and conducted all experiments. MP and AC contributed to device development. SGM and BYC coordinated efforts. KS, MSM, and BYC analyzed data and prepared the manuscript and assembly guides. KS, MP, and AC conducted their work as Penn iGEM 2017.

## Acknowledgments

The authors thank Chris Fang-Yen for helpful discussion on optical train design, Thomas Gilgenast for helpful discussion on API design, Erin Berlew for EL222 recombinant protein production, and Kevin Gardner for the EL222 plasmid. The Ocean Optic logo was used with permission from the company; the OSP is not a product of the company and its development was not sponsored by the company. This work was funded by the George H. Stephenson Foundation, the University of Pennsylvania Department of Bioengineering, the University of Pennsylvania Office of Vice Provost for Research (VPR), the National Science Foundation (NSF, CAREER Award MCB 652003), and the National Institutes of Health (NIH, R01NS101106). MSM was partially supported by an NSF GRFP Award.

## Abbreviations

AHL: Acyl-homoserine lactone
API: Application Programming Interface
CCD: Charge-coupled device
EL222: Light-activated DNA-binding protein EL222
GFP: Green fluorescent protein
GUI: Graphical User Interface
LED: Light-emitting diode
LOV: Light-oxygen-voltage
luxR: Transcriptional activator protein luxR
OD: Optical density
PMT: Photomultiplier tube
RBS: Ribosome binding site
SMA: Sub-miniature version A interface

